# Evolutionary genomics based on PacBio HiFi long-read sequencing data reveals the importance of structural variants in shaping population-specific differences between Chinese and Indian rhesus macaques (*Macaca mulatta*)

**DOI:** 10.64898/2026.05.27.728199

**Authors:** Takahiro Maruki, Cyril J. Versoza, Jeffrey D. Jensen, Susanne P. Pfeifer

## Abstract

Rhesus macaques (*Macaca mulatta*) are the most widely used non-human primate model for translational research relevant to human health and disease. Although several genetically distinct populations have been recognized across the species’ extensive habitat range in Asia, the majority of biomedical studies in the United States and abroad focuses on individuals of either Chinese or Indian descent. Notably, phenotypic differences exist between these two populations which can influence biomedical research outcomes; however, the genetic basis and molecular mechanisms underlying these differences are generally not well understood. Based on novel PacBio HiFi long-read sequencing data from 20 rhesus macaques — ten of Chinese origin and ten of Indian origin — we here characterize the genome-wide landscape of structural variation in these two biomedically-relevant populations. Our results highlight differences in the structural variant landscape affecting genes involved in neural communication and signaling pathways, in line with the known differences in temperament between the two populations. Furthermore, while the majority of discovered structural variants were located in intergenic and non-coding regions of the genome, 15 of the discovered population-specific structural variants were predicted to exhibit a high functional effect on genes associated with human disease, indicating that they may play an important role in shaping the differences in disease susceptibility between the populations. Taken together, by providing detailed insights into population-specific structural variation, this genomic resource will aid the design and interpretation of future studies aiming to link genotype, phenotype, and fitness in the context of human health and disease, and facilitate broader comparative analyses of structural variation as a force shaping genome evolution across primates.

## INTRODUCTION

Rhesus macaques (*Macaca mulatta*) are one of the most important models for biomedical research, owing to their similarities to humans in terms of behavior, cognition, development, and physiology as well as susceptibility to infectious agents (Rhesus Macaque Genome Sequencing and Analysis Consortium et al. 2007; and see the reviews of Cooper et al. 2022; Vallender et al. 2023). Specifically, rhesus macaques are widely used as translational models for neurodevelopmental disorders (including schizophrenia; Simen et al. 2009), neurodegenerative diseases (including autism spectrum disorder and Alzheimer’s disease; Bauman and Schumann 2018; Stonebarger et al. 2021), metabolic diseases (including cardiovascular disease and type-2 diabetes; Havel et al. 2017), and substance use disorders (Banks et al. 2017). Additionally, the species serves as a critical model for the study of infection dynamics for a variety of pathogens with diverse etiologies and relevance to human health and disease (including emerging, or re-emerging, diseases such as Ebola, HIV, SARS-CoV2, tuberculosis, and Zika; Dudley et al. 2016; McMahan et al. 2021; and see the review of Gardner and Luciw 2008).

Rhesus macaques are geographically widespread, with a natural habitat extending from the Hindu Kush of Afghanistan and Pakistan, across the Himalayas (India, Nepal, and Bhutan), into the lowlands and tropical forests of mainland Southeast Asia (Myanmar, Thailand, Laos, and Vietnam), and reaching eastward into southern China — habitats that encompass a diverse topology, with elevations ranging from sea level to 4,000 m (Groves 2001). In addition to this geographic diversity, rhesus macaques also display a remarkable heterogeneity at both the genetical and morphological levels, and multiple distinct populations have been recognized to date (Groves 2001). Among the most important from a biomedical perspective are the populations of Chinese and Indian descent as individuals from these populations served as founders for many primate research centers across the globe (with most individuals housed in the United States originating from India and those of Chinese descent being largely used within their home country; Rogers 2022). From a phenotypic perspective, Chinese rhesus macaques tend to be larger and exhibit a more pronounced sexual dimorphism than Indian rhesus macaques (Clarke and O’Neil 1999); additionally, Chinese individuals tend to be more aggressive (Champoux et al. 1994,1997; Jiang et al. 2013). Previous studies have also described genetic variation linked to phenotypes that can influence biomedical outcomes, including differences in blood chemistry (Champoux et al. 1996), major histocompatibility complex alleles (Viray et al. 2001; Doxiadis et al. 2003), as well as infectious disease dynamics, with individuals of Chinese descent, for example, being more resistant to simian immunodeficiency viruses than individuals of Indian descent (Joag et al. 1994; Ling et al. 2002; Trichel et al. 2002; and see the review of Zhou et al. 2013). Gaining a better understanding of the genetic drivers of these observed phenotypic differences is thus critically important to improve both the usage of rhesus macaques as biomedical models for human health and disease as well as to inform evolutionary research.

Compared to humans, rhesus macaques are characterized by a relatively high level of genetic diversity (Warren et al. 2020) and substantial structuring exists between populations of Chinese and Indian descent (*F_ST_* ∼ 0.15), with limited gene flow since their split during the Early Pleistocene (∼1.5 million years ago; Hernandez et al. 2007; Heenkenda et al. 2026). Moreover, while the Chinese population expanded substantially after the population split, the Indian population experienced a relatively strong bottleneck, resulting in an effective population size that is more than an order of magnitude lower than that of the Chinese population (∼14,000 vs ∼220,000; Heenkenda et al. 2026). Additionally, notable differences exist in the landscape of recombination between the two populations, particularly at the finer scales (Spatola et al. 2026). Given these differences and their expected impact on translational research, the genetic variation carried by these two populations has thus been the focus of extensive genomics efforts and several databases now exist that offer insights into the landscapes of single nucleotide variation in the species (Zhang et al. 2013; Bimber et al. 2019; Warren et al. 2020). By contrast, in spite of the often profound effects of structural variants (i.e., deletions, duplications, insertions, and inversions > 50 bp) on gene structure, gene expression, and regulation — as well as their frequent involvement in both common, rare, and complex diseases (see the reviews of Hurles et al. 2008; Weischenfeldt et al. 2013; Lupski 2015) — high-quality population-scale genomic resources for structural variation remain sparse due to the inherent difficulties of (the still dominant) short-read sequencing to comprehensively characterize structural genomic changes (though see Ray et al. 2025; Zhang, Xu et al. 2025). As a result, insights into how structural variation shapes population-specific genomic landscapes remain largely elusive.

Taking advantage of newly generated whole-genome PacBio high-fidelity (HiFi) long-read sequencing data from 20 unrelated rhesus macaques — ten of Chinese descent and ten of Indian descent, we here characterize the genomic landscape of structural variation in the species and further infer how population-private structural variants may shape the phenotypic differences observed between these two biomedically-relevant populations. By providing novel insights into population-specific structural variation, this genomic resource will thus not only allow for an improved design and interpretation of future genotype to phenotypic to fitness studies performed in the colonies, but it will also enable more comprehensive comparisons of structural variation as a driver of genome evolution across the primate lineage.

## RESULTS & DISCUSSION

### Structural variant catalogue of rhesus macaques of Chinese and Indian descent

PacBio HiFi long-reads were generated for 20 unrelated rhesus macaques (*Macaca mulatta*) — ten individuals of Chinese descent and ten individuals of Indian descent (Figure 1a) — by whole-genome sequencing their genomic DNA on the Revio platform. Sequencing coverage was similar for individuals originating from the Chinese population and those originating from the Indian population — with an average depth of 11.7× (range: 6.3–16.2×) and 13.3× (range: 10.5–16.7×), respectively (Wilcoxon rank-sum test, *p*-value = 0.325; Figure 1b; and see Supplementary Table S1) — but with significantly longer read lengths in the Chinese population (21.6 kb vs 18.4 kb; Wilcoxon rank-sum test, *p*-value = 0.007; Figure 1c). PacBio HiFi reads were quality-controlled with LongQC (Fukasawa et al. 2020) and HiFiAdapterFilt (Sim et al. 2022), confirming the absence of contamination and adapter sequences, before aligning them to the rhesus macaque reference assembly, Mmul_10 (Warren et al. 2020), using minimap2 (Li 2018, 2021). Although the reference assembly was built from a female of Indian-origin, reads from both populations aligned equally well to this reference assembly (with a mean of >99.99%; Wilcoxon rank-sum test, *p*-value = 0.091; Supplementary Table S2); however, alignment identity was higher for the Indian population (98.1 in individuals of Indian origin vs 97.7 in individuals of Chinese origin on average; Wilcoxon rank-sum test, *p*-value = 1.81 × 10^-4^), as expected. In contrast, the mean mapping quality was higher in the Chinese population (with a mean of 39.4 compared to 38.2 in the Indian population; Wilcoxon rank-sum test, *p*-value = 2.17 × 10^-5^), in agreement with the overall longer read lengths. Structural variants were called from these long-read alignments using Sniffles2 (Sedlazeck et al. 2018; Smolka et al. 2024), and genotypes of insertions and deletions with a length of ≤ 10 kb were further refined using the *k*-mer genotyper kanpig (English et al. 2025) to improve accuracy. In total, 397,719 structural variants were discovered in the 20 individuals of this study (Supplementary Table S3).

**Figure 1.**
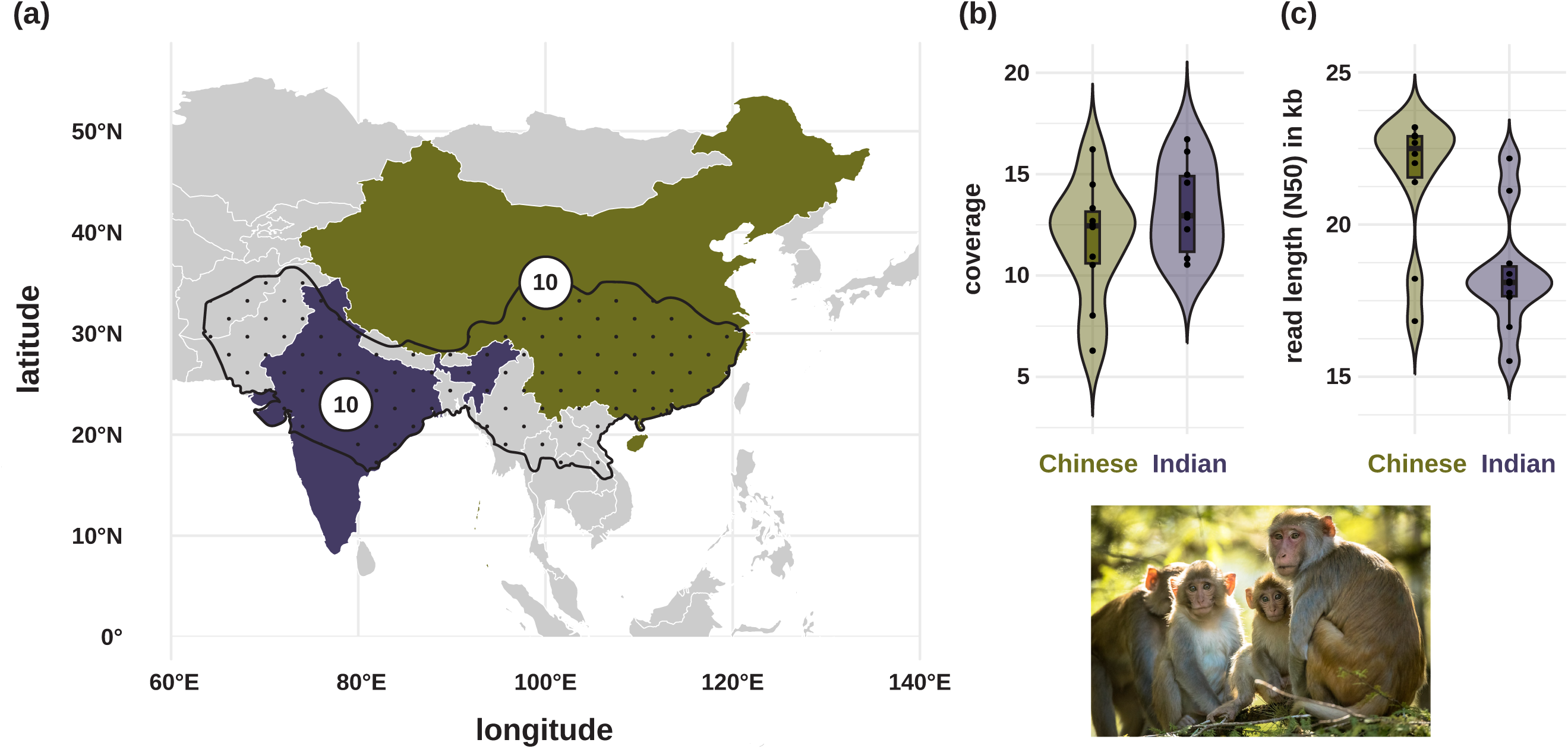
**(a)** The geographic distribution of rhesus macaques (*Macaca mulatta*) ranges from Afghanistan and Pakistan across the Indian subcontinent into Myanmar and across China to the southeastern coast (species range shown as the dotted area, relative to current national borders given by white lines). For this study, 10 individuals each of Chinese (green) and Indian (purple) descent were sampled and long-read sequenced using PacBio HiFi sequencing. Range information was obtained from the International Union for the Conservation of Nature. **(b)** Average genome-wide coverage per individual. **(c)** Average read length (N50) per individual. Picture credit: "71279" by Rachel Simmons (CC BY-NC-ND 2.0).

### The landscape of structural variation in rhesus macaques of Chinese and Indian descent

Substantial differentiation was observed between the two rhesus macaque populations (*F_ST_* = 0.158), with individuals of Chinese descent exhibiting a higher structural variant diversity compared to individuals of Indian descent, harboring ∼93k (range: 84k–97k) and ∼76k (range: 72k–77k) variants per individual on average, respectively (Wilcoxon rank-sum test, *p*-value = 1.08 × 10^-5^; Supplementary Table S3). Although the landscape of structural variation in both populations was largely governed by insertions (59.52% and 61.20% per individual on average in the Chinese and Indian populations, respectively) and deletions (40.38% and 38.70%) — with additional minor contributions of inversions (0.08% and 0.09%) and duplications (0.01% and 0.02%) — the two populations differed significantly in their overall structural variant composition (*F*_1,18_ = 102.29, *R*^2^ = 0.850, *p*-value = 0.001; Figure 2). Moreover, a larger proportion of variants were private to the Chinese population compared to the Indian population, as expected owing to the larger effective population size in the former (Heenkenda et al. 2026) and consistent with previous studies focused on single nucleotide variants (Ferguson et al. 2007; Hernandez et al. 2007). Among the different classes of structural variants, deletions, inversions, and insertions exhibited the highest proportion of sharing (43%, 39%, and 29%, respectively), suggesting that these variants either arose through recurrent mutation in regions prone to structural rearrangements or predate the population split. In contrast, duplications were predominantly population-specific (with only 18% of variants shared between the two populations), consistent with a greater contribution of more recent, lineage-specific structural changes. In general, population-specific structural variants tended to segregate at low frequencies — with slightly higher minor allele frequencies in the Indian population than in the Chinese population — and structural variants shared between the two populations were at intermediate to high allele frequencies.

**Figure 2.**
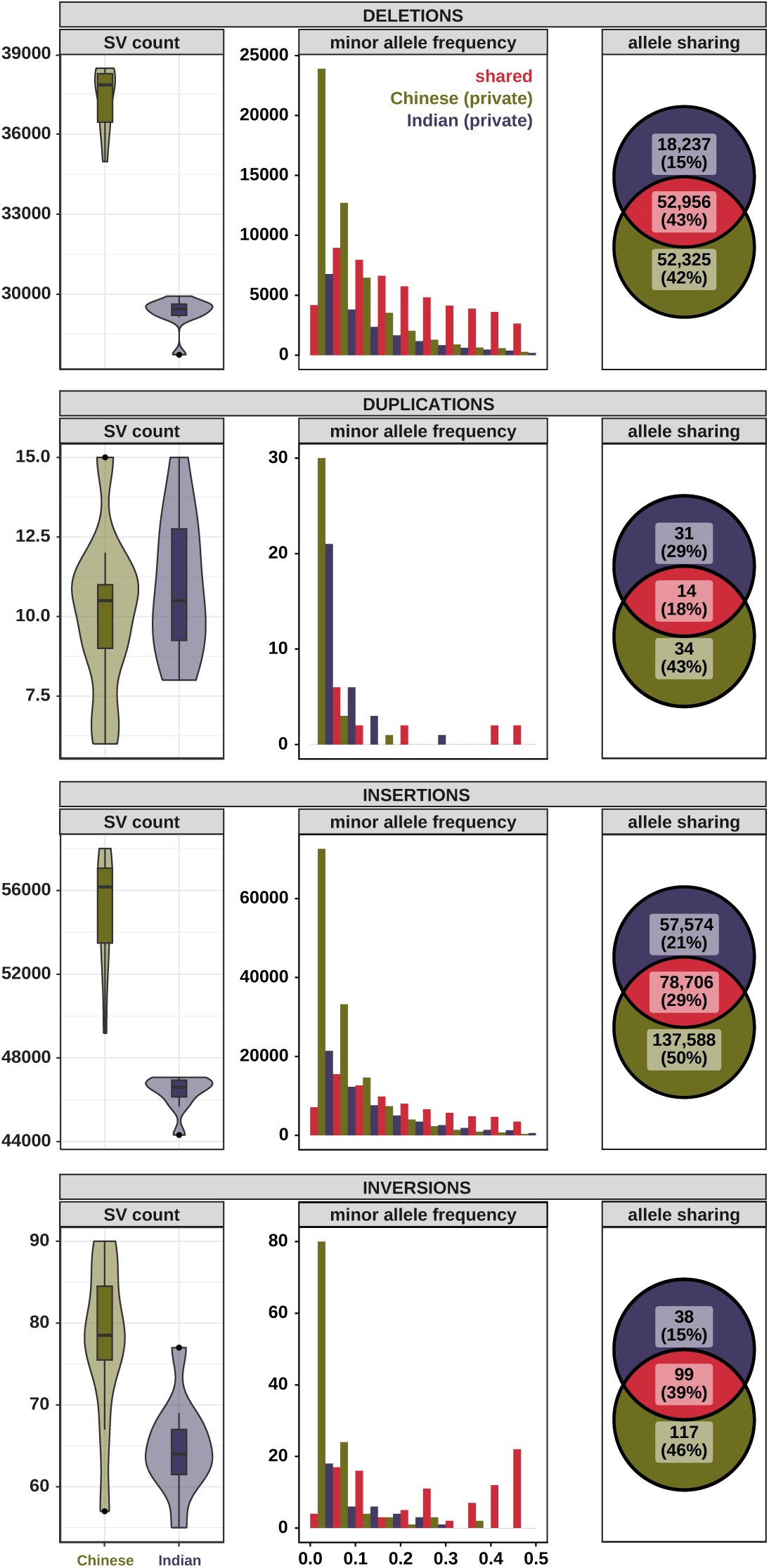
Landscape of structural variation in rhesus macaques of Chinese and Indian descent. **(Left)** Number of structural variants (SVs) per individual of Chinese and Indian descent (shown in green and purple, respectively). (**Middle**) Distribution of minor allele frequencies of structural variants that are private to the Chinese population (shown in green), private to the Indian population (purple), and shared between the two populations (pink). (**Right**) Number (proportion) of alleles private to the Chinese population (shown in green), private to the Indian population (purple), and shared between the two populations (pink).

Statistically significant differences between the two rhesus macaque populations were also observed with regards to the length of the structural variants (Supplementary Figure S1), with deletions and insertions private to the Chinese population being longer than those private to the Indian population (Wilcoxon rank-sum tests, *p*-value_DEL_ = 2.20 × 10^-2^ and *p*-value_INS_ = 2.35 × 10^-35^), though these differences were small in magnitude (Chinese_DEL_: 136 ± 890 bp vs Indian_DEL_: 131 ± 950 bp; Chinese_INS_: 308 ± 1,050 bp vs Indian_INS_: 304 ± 1,080 bp) and are thus unlikely to be biologically meaningful. A more pronounced difference was however observed between population-private and shared deletions, with the latter being substantially longer (shared_DEL_: 215 ± 1,009 bp; Wilcoxon rank-sum test, *p*-value = 9.39 × 10^-167^).

The distribution of structural variants across the rhesus macaque genome was highly heterogeneous. At the broad scale, structural variants were depleted on the longest chromosomes whereas enrichments were observed on the shorter chromosomes (Figure 3a). GC-content — itself strongly correlated with repeat-content (*r* = 0.66) — was one of the strongest predictors of structural variant density at the chromosomal scale (Supplementary Figure S2), in agreement with previous studies in humans showing that GC-content impacts structural variation, particularly that mediated by mobile elements (Audano et al. 2019; Lim et al. 2025). Notably, the chromosome with the highest GC-content, chromosome 19 (48.34% compared to the autosomal mean of 41.48%), exhibited the highest enrichment of structural variants of each class (Figure 3a), consistent with the hypermutable nature and accelerated nucleotide evolution previously reported for this chromosome across the primate clade (Harris et al. 2020). At the fine (1 Mb) scale, structural variants clustered in sub-telomeric regions and in close proximity to centromeres (Figure 3b), genomic regions prone to double-strand breaks due to their high density of repetitive DNA (Audano et al. 2019). Additionally, an enrichment of structural variants was observed within the major histocompatibility complex (located on chromosome 4) in both populations, in agreement with previous work in various primate lineages (Mao et al. 2024).

**Figure 3.**
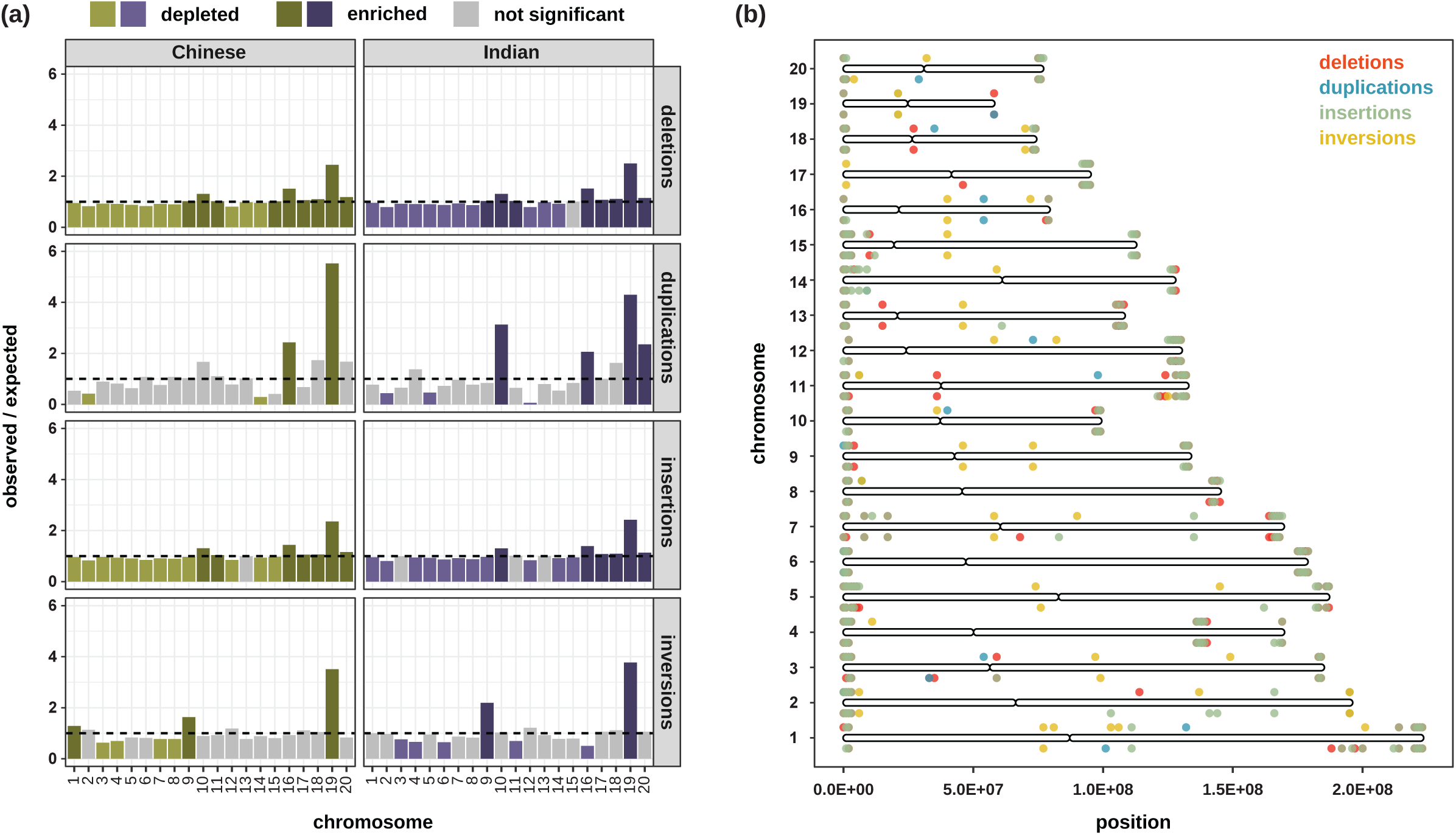
Distribution of structural variants along the rhesus macaque genome. **(a)** Enrichment (dark colors) and depletion (light colors) of structural variants per chromosome in populations of Chinese (shown in green) and Indian (purple) descent. The horizontal dashed line indicates the expectation under a uniform distribution (i.e., an observed vs expected ratio of 1). Enrichments / depletions of structural variants on chromosomes shaded in gray are not statistically significant. (b) Fine-scale structural variant landscapes plotted in 1 Mb bins along each chromosome. Structural variant hotspots (*z* ≥ 2) are shown as colored points (with deletions shown in red, duplications in teal, insertions in sage green, and inversions in yellow) above (Chinese) and below (Indian) the chromosome ideograms.

The vast majority of structural variants were located in intergenic and non-coding regions of the genome (between 73.7% and 91.2%, depending on structural variant class and population; Figure 4), as expected from the genome-wide composition (Warren et al. 2020). No statistically significant differences were observed in the distributions of deletions and insertions across functional categories (χ^2^_del_ = 8.71, df = 5, *p*-value = 0.1213; χ^2^_ins_ = 5.20, df = 4, *p*-value = 0.2678); however, a larger proportion of protein-coding changes was observed for both duplications and inversions in the Indian population compared to the Chinese population (16.1% vs 2.9% for duplications and 26.3% vs 14.5% for inversions) as might be expected from its smaller effective population size and thus reduced efficacy of purifying selection (Heenkenda et al. 2026), though the number of observations was small. Accordingly, most structural variants were predicted to be modifiers with little to no effect (Supplementary Figure S3), as anticipated given that only a small fraction (∼1–2%) of the rhesus macaque genome is coding (Warren et al. 2020); nevertheless, a small proportion (<2%) of deletions and insertions were predicted to be of high functional impact.

**Figure 4.**
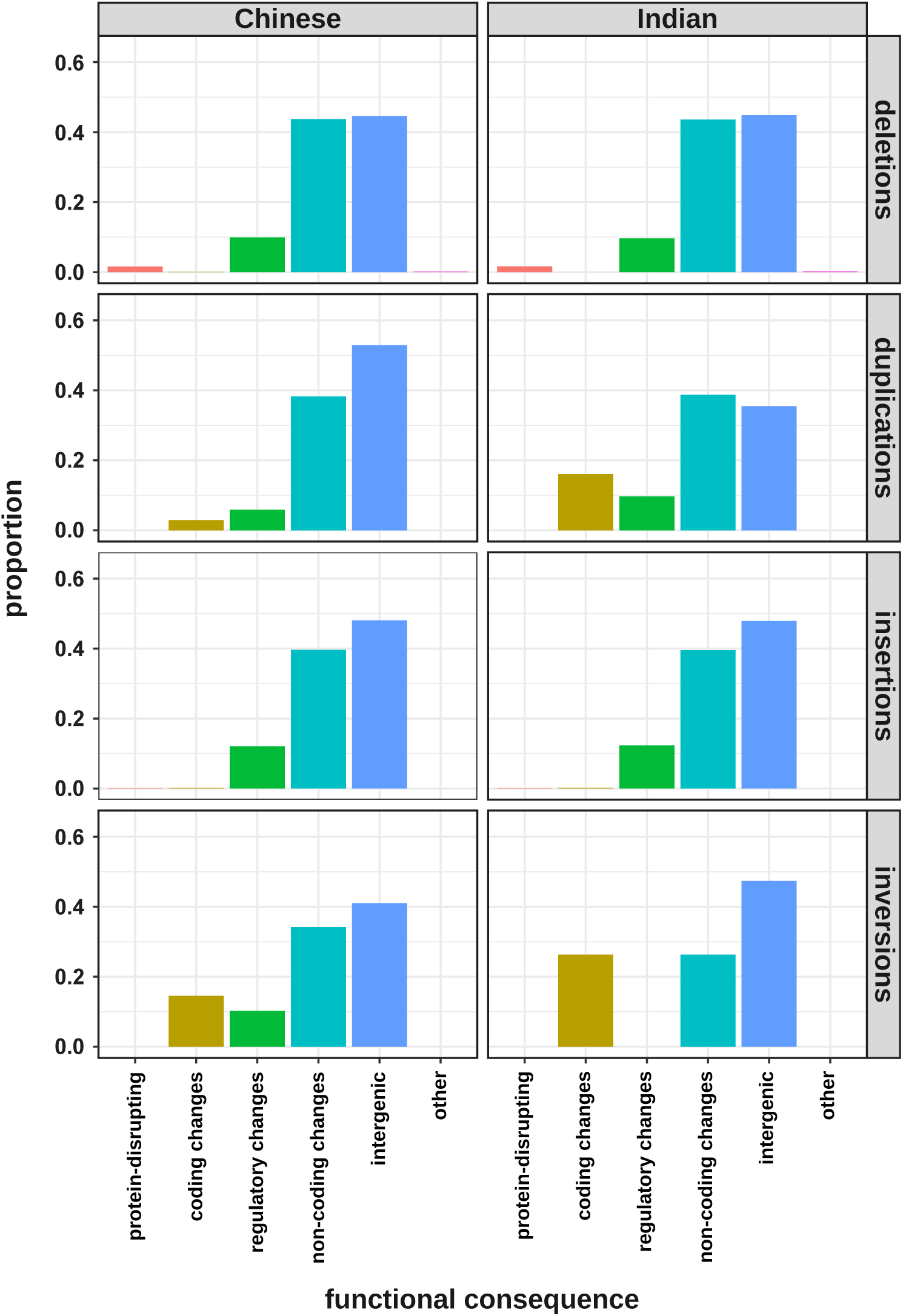
Distribution of population-specific structural variants across six functional categories: protein-disrupting (consisting of frameshift variants, splice acceptor/donor variants, lost start/stop codons, transcript ablations, and feature truncations), coding changes (codon sequence/transcript variants), regulatory changes (downstream/upstream gene variants, 3’/5’ UTR variants, splice region variants, and feature elongations), non-coding transcript changes (intron variants, non-coding transcript variants, transcript amplification, and miRNA variants), intergenic changes (intergenic variants), and other (all remaining annotations).

### The landscape of mobile elements in rhesus macaques of Chinese and Indian descent

Mobile elements — an important source of structural variation — in rhesus macaques of Chinese and Indian descent were identified from the long-read data using rMETL (Jiang et al. 2019), based on known repetitive DNA elements in primates available in the Dfam database (Storer et al. 2021). Similar to other structural variants, Chinese individuals harbored a significantly larger number of segregating mobile elements than Indian individuals (Wilcoxon rank-sum test, *p*-value = 7.58 × 10^-5^; Figure 5a). The landscape of recently or currently active mobile elements in both populations was dominated by long interspersed nuclear elements (LINEs) and short interspersed nuclear elements (SINEs). Specifically, consistent with previous work in rhesus macaques and other primates (Han et al. 2007; Konkel et al. 2010; Hormozdiari et al. 2013; Tang and Liang 2019; Warren et al. 2020; Chabukswar et al. 2022; Williams et al. 2024), truncated autonomous LINE-1 (L1) retrotransposons (mean lengths: 1.3 and 1.5 kbs in the Chinese and Indian population, respectively) and non-autonomous SINE/*Alu* elements (mean lengths: 324 and 338 bps) were the most abundant, with the former contributing 70.7% and 68.6% — a proportion similar to that previously observed in humans (71%; Lam et al. 2010). Additionally, minor contributions of HERV-K(HML-2)-like elements were observed in both populations (mean lengths: 4.5 and 5.5 kbs; Figure 5b). Despite these similarities, the relative distribution of mobile element classes differed significantly between the two populations (χ^2^ = 84.87, df = 2, *p*-value < 2.20 × 10^-16^; Figure 5c). The rates of L1 insertions and *Alu* retrotransposition are known to vary widely between different primate species, with *Alu* retrotransposition being the most prevalent in humans, chimpanzees, bonobos, and gorillas, whereas a higher accumulation of L1 insertions has been found in orangutans as well as the common ancestor of humans and non-human apes (Hormozdiari et al. 2013; Yoo et al. 2025). Being largely selectively neutral and distributed randomly throughout the genome, fixed *Alu* insertions have indeed proven useful to study phylogenetic relationships (see the review of Xing et al. 2007); similarly, segregating loci can aid in our understanding of the structuring between populations. A principal component analysis of segregating *Alu* elements showed a strong structuring between Chinese and Indian populations, in agreement with both the *F_ST_* value observed from genome-wide structural variation and previous work based on single nucleotide polymorphisms (Hernandez et al. 2007; Heenkenda et al. 2026).

**Figure 5.**
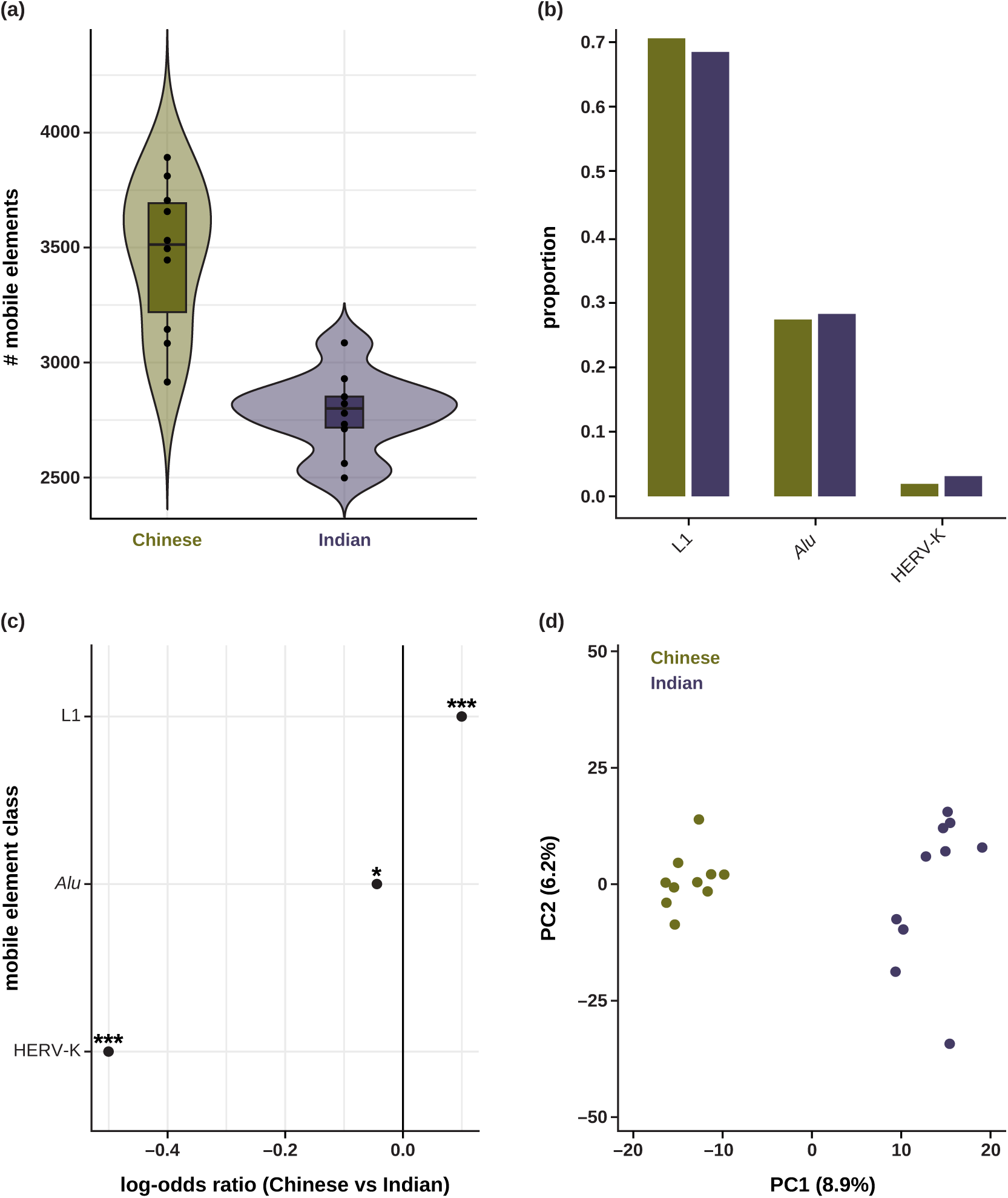
Landscape of mobile elements in rhesus macaques of Chinese and Indian descent. **(a)** Number of mobile elements per individual of Chinese and Indian descent (shown in green and purple, respectively). **(b)** Relative contribution of mobile elements with current or recent activity in primates (*Alu*, L1, and HERV-K(HML-2)-like elements) per population. **(c)** Log-odds ratio plot of the relative contribution of mobile elements with current or recent activity in primates (*Alu*, L1, and HERV-K(HML-2)-like elements), with 0 (solid black line) indicating no difference between the two populations, positive values indicating an enrichment in the Chinese population, and negative values indicating an enrichment in the Indian population. * and *** indicate a *p*-value < 0.05 and < 0.001, respectively. **(d)** Principal component analysis of *Alu* elements segregating in the Chinese and Indian populations.

### Mutational mechanisms underlying structural variant formation

Gaining a better understanding of the mutational mechanisms underlying structural variant formation is important to inform evolutionary and population genomics research. Different mechanisms leave distinct signatures in the genome, with non-homologous end-joining (NHEJ) resulting in blunt joins with an absence of homology at the breakpoint junctions, microhomology-mediated end-joining (MMEJ) exhibiting short (usually around 1–15 bps) microhomology, and non-allelic homologous recombination (NAHR) being characterized by long (often ≥ 100 bps) homologous sequences at the flanking breakpoints (Currall et al. 2013). Long-read sequencing allows for the characterization of the precise breakpoints of structural variants, and patterns of microhomology in genomic regions flanking these breakpoints can thus be leveraged to infer the putative mutational mechanisms underlying the formation of structural variants (see the reviews of Ottaviani et al. 2014; Weckselblatt and Rudd 2015). A substantial proportion (26.3%) of the structural variation in the two rhesus macaque populations was driven by mobile elements, consistent with previous observations in humans (20.6 % in Kidd et al. 2010; 21.3% in Lam et al. 2010). Out of the remainder, NHEJ played the most important role in the formation of structural variation (68.0%), as expected given the prevalence of this double-strand break repair mechanism in mammals (Mao et al. 2008) and its activity throughout all cell cycle phases (Chang et al. 2017). Primarily active during the S and G2 phases of the cell cycle (see the review of Sfeir et al. 2024), MMEJ created around a third (30.3%) of non-mobile element structural variants observed in the rhesus macaque populations whereas NAHR, which is operating alongside MMEJ and is highly active during meiosis, contributed only a minor fraction (1.7%).

### Phenotypic traits associated with population-private structural variation

To investigate structural variation potentially shaping the phenotypic differences between the rhesus macaque populations of Chinese and Indian descent utilized in biomedical research, we focused on variation private to a single population. At a statistical significance level of 5%, more private structural variants were observed in the Indian population than in the Chinese population (3,367 vs 2,714), consistent with an excess of intermediate- and high-frequency variants owing to the demographic history of the Indian population (Heenkenda et al. 2026). Notably, biological processes related to neuronal signaling, including modulation of chemical synaptic transmission (3.80-fold enrichment, FDR = 7.3 × 10^-4^), intracellular signal transduction (1.98-fold, FDR = 3.7 × 10^-3^), and protein phosphorylation (1.90-fold, FDR = 4.5 × 10^-3^) were overrepresented among the structural variation private to the Indian population (Table 1; and see Supplementary Table S4 for a complete list of gene ontology [GO] terms enriched within genes intersecting the Indian-private structural variants). Consistently, enriched cellular components included dendric spine (2.88-fold, FDR = 2.4 × 10^-2^E) and postsynaptic membrane (2.26-fold, FDR = 2.7 × 10^-2^), whereas molecular functions were dominated by ion channel activity involved in the regulation of pre- and postsynaptic membrane potential (6.92-fold, FDR = 1.4 × 10^-2^ and 4.92-fold, FDR = 2.4 × 10^-3^, respectively). Similarly, Chinese-private structural variants were enriched in genes associated with microtubules (2.16-fold, FDR = 4.4 × 10^-2^; Table 1; and see Supplementary Table S5), which play an important role in signal transduction and the nervous system (see the review of Jang et al. 2024). Taken together, these results thus highlight important differences in the structural variant landscape affecting genes involved in neural communication and signaling pathways, in line with the known differences in temperament and disease susceptibility between the two populations (Champoux et al. 1997; Trichel et al. 2002).

**Table 1.**
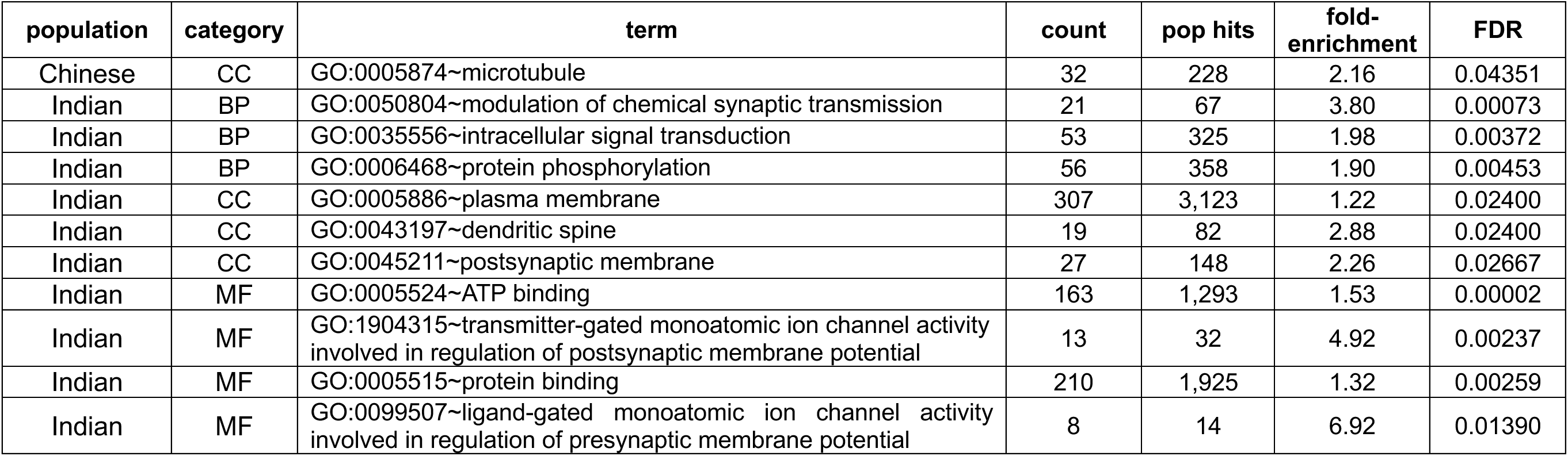
Gene ontology (GO) terms enriched in genes intersecting significant population-private structural variants. BP, CC, and MF denote the biological process, cellular component, and molecular function, respectively. Count and pop hits denote the numbers of genes associated with each term in the significant population-private structural variant catalogues and in the genomic background, respectively. FDR denotes the false discovery rate.

| population | category | term | count | pop hits | fold-enrichment | FDR |
| --- | --- | --- | --- | --- | --- | --- |
| Chinese | CC | GO:0005874~microtubule | 32 | 228 | 2.16 | 0.04351 |
| Indian | BP | GO:0050804~modulation of chemical synaptic transmission | 21 | 67 | 3.80 | 0.00073 |
| Indian | BP | GO:0035556~intracellular signal transduction | 53 | 325 | 1.98 | 0.00372 |
| Indian | BP | GO:0006468~protein phosphorylation | 56 | 358 | 1.90 | 0.00453 |
| Indian | CC | GO:0005886~plasma membrane | 307 | 3,123 | 1.22 | 0.02400 |
| Indian | CC | GO:0043197~dendritic spine | 19 | 82 | 2.88 | 0.02400 |
| Indian | CC | GO:0045211~postsynaptic membrane | 27 | 148 | 2.26 | 0.02667 |
| Indian | MF | GO:0005524~ATP binding | 163 | 1,293 | 1.53 | 0.00002 |
| Indian | MF | GO:1904315~transmitter-gated monoatomic ion channel activity involved in regulation of postsynaptic membrane potential | 13 | 32 | 4.92 | 0.00237 |
| Indian | MF | GO:0005515~protein binding | 210 | 1,925 | 1.32 | 0.00259 |
| Indian | MF | GO:0099507~ligand-gated monoatomic ion channel activity involved in regulation of presynaptic membrane potential | 8 | 14 | 6.92 | 0.01390 |

Among the significant population-private structural variants, 15 insertions were predicted to exhibit a high functional effect on genes associated with diseases in humans (Table 2). Of particular note for the population of Chinese descent, an insertion was predicted to affect *DISC1*, a gene essential for neurodevelopmental processes including neuronal proliferation and migration as well as intracellular signalling. In humans, mutations in *DISC1* — both common non-synonymous and rare missense single nucleotide polymorphisms (see Soares et al. 2011 and references therein) as well as structural variants (Song et al. 2008) — have been linked to a variety of mental illnesses, including a susceptibility to schizophrenia. In contrast to rodents, the nucleotide sequence and amino acid structure of *DISC1* are highly conserved between macaques and humans (Bord et al. 2006), suggesting a conserved role in neurodevelopment. Together with profound similarities in brain development and function — particularly in terms of complex cortical expansion (with structural abnormalities in the prefrontal cortex observed in schizophrenic individuals) — macaques thus serve as an important translational model system (see the reviews of Capitanio and Emborg 2008; Simen et al. 2009; Cheng et al. 2025). Of note in this regard, employing a reverse genomics approach, Zhang, Chen et al. (2025) recently identified a missense mutation in *DISC1* (p.Arg517Trp) segregating in a cohort of >900 captive rhesus macaques of Chinese descent that resulted in phenotypic alterations in both cortical architecture and neurological function. Additionally, research in closely related cynomolgus macaques (*M. fascicularis*) demonstrated that CRISPR-Cas9-introduced mutations in *DISC1* can cause complex behavioral phenotypes resembling those observed for human psychiatric conditions (Zhou et al. 2026). Relatedly, *PDE10A*, which encodes an enzyme expressed in the brain’s striatum, has been implicated in schizophrenia as well as other neuropsychiatric disorders and serves as a promising target in pharmacological research (see the reviews of Menniti et al. 2006; Rautela et al. 2026). Given the previous usage of rhesus macaques as a model system for the study of the behavioral effects of *PDE10A* inhibitors (Smith et al. 2013; Uthayathas et al. 2014), the observation of a structural variant predicted to affect this gene is noteworthy.

**Table 2.**
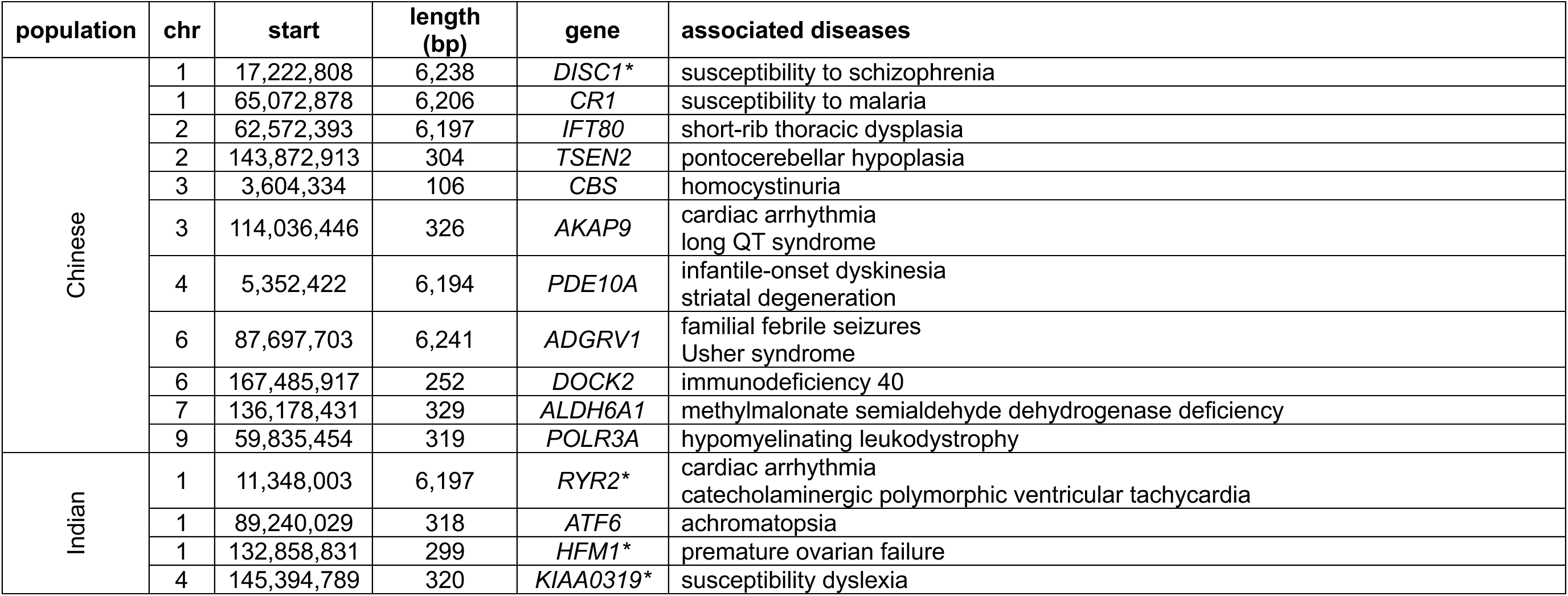
Population-specific structural variants predicted to be of high effect on genes associated with human disease. Genes associated with population-specific GO term enrichments (Table 1) are highlighted by a star (*).

With regards to infectious disease, an insertion was observed to be segregating in *CR1*, one of the most structurally complex genes involved in both innate and adaptive immunity. *CR1* comprises long stretches of homologous repeats, making it prone to structural variation, which in turn results in different protein isoforms and expression levels on erythrocytes (Khera and Das 2009). In humans, mutations in this gene have been associated with malaria susceptibility (see the reviews of Kwiatkowski et al. 2005; Rowe et al. 2009; Zhang, Wu et al. 2025 and references therein), with signatures of positive selection observed in populations inhabiting malaria-endemic regions (Kosoy et al. 2011; Lorenzini et al. 2023). As a natural host for several *Plasmodium* species that cause malaria (Schmidt et al. 1961), rhesus macaques are an important model to improve our understanding of host-pathogen dynamics and thus the structural variation identified here may serve as an interesting candidate for future malaria studies conducted in the primate research colonies.

In addition to neurobiological and immunological research, rhesus macaques also serve as a translational model for metabolic research, including studies focused on alcohol substance disorders (see the review of Grant and Bennett 2003). The identification of an insertion predicted to affect *ALDH6A1*, a gene which encodes an enzyme involved in both the detoxification of aldehydes and energy metabolism in the liver, is thus of interest.

From a physiological point of view, rhesus macaques have well-characterized cardiac arrhythmia phenotypes, including cardiomyopathy-associated sudden death (Kanthaswamy et al. 2014; Ueda et al. 2021; Rivas et al. 2024). In this context, the observation of a population-private structural variant in individuals of Indian descent predicted to impact *RYR2*, a gene encoding a cardiac ryanodine receptor necessary for the regulation of calcium during heart contractions, is intriguing given that mutations in this gene have been associated with a variety of cardiac arrhythmias in humans (see the review of Lv et al. 2025); similarly, a population-private structural variant segregating in the Chinese population was predicted to affect *AKAP9*, mutations of which have been implicated in cardiovascular diseases in humans (see the review of Zhang, Zhu et al. 2025).

Lastly, from an evolutionary standpoint, the structural variant discovered in *HFM1*, a gene necessary for crossover formation and chromosome synapsis during meiosis (Guiraldelli et al. 2013), is notable for its relationship to primary ovarian insufficiency, infertility, or other reproductive issues observed in humans (Pu et al. 2016; Tang et al. 2021).

Taken together, the observed differences between the structural variant landscapes of Chinese and Indian rhesus macaques thus underscore the need to carefully select the population most suitable for a specific research endeavor, and the novel genomic resources developed here are expected to aid both the design and interpretation of future biomedical and evolutionary studies aiming to link genotype, phenotype, and fitness in this species.

## MATERIALS AND METHODS

### Animal subjects

Rhesus macaques were housed in indoor or outdoor social housing at the Oregon National Primate Research Center (ONPRC). All husbandry practices conducted are performed in accordance with federal guidelines and regulations as stated in the National Institutes of Health Guide for the Care of and Use of Laboratory Animals. ONPRC is accredited by the Association for Assessment and Accreditation of Laboratory Animal Care, International. Buffy coat samples were previously collected and stored under Oregon Health and Science University (OHSU) IACUC protocol #IP00000367.

### Samples and PacBio HiFi sequencing

Previously collected high molecular weight DNA was obtained from 20 unrelated, captive rhesus macaques (*Macaca mulatta*) — 10 individuals of Chinese-descent and 10 individuals of Indian-descent — housed at the Oregon National Research Center. DNA from each individual was sheared to a 10–20 kb size range using a Megaruptor 3 (Diagenode, Liège, Belgium). Afterward, sheared DNA was purified using SMRTbell cleanup beads and sequencing libraries were constructed using the SMRTbell Prep kit 3.0, following manufacturer protocols. Sequencing libraries were size selected on a Pippin HT (Sage Science, Beverly, MA, USA) using S1 markers with a 10–25 kb size selection. Final libraries were quantified with a Qubit HS kit (Invitrogen, Carlsbad, CA, USA) and size checked on a Femto Pulse System (Agilent, Santa Clara, CA, USA) before preparing them for sequencing using the PacBio Sequel II Sequencing kit 3.1 for HiFi libraries, loading them on Revio SMRT cells, and sequencing them in the CCS mode for 24 hours. Sample information is provided in Supplementary Table S1.

### Long-read pre-processing and alignment

The quality of the raw long-read data was examined using LongQC v.1.2.0c (Fukasawa et al. 2020), specifying the PacBio HiFi preset (*-x* pb-hifi), and reads with remnant adapter sequences were removed using HiFiAdapterFilt v.3.0.0 (Sim et al. 2022). Quality-controlled reads for each individual were aligned to the rhesus macaque reference assembly, Mmul_10 (GenBank accession number: GCA_003339765.3; Warren et al. 2020), using minimap2 v.2.30 (Li 2018, 2021), specifying the PacBio HiFi preset (*-ax* map-hifi) and adding read group information (*-R*). Summary statistics of the autosomal read alignments for each individual, calculated using SAMtools *flagstat* v.1.21 (Li et al. 2009), are provided in Supplementary Table S2. Population-specific differences for each sequencing metric (percent aligned reads, alignment identity, and mean mapping quality) were assessed using two-sided Wilcoxon rank-sum tests in R v.4.4.2 (R Core Team 2026).

### Structural variant calling and genotyping

For each individual, autosomal candidate structural variants were called from the long-read alignments using Sniffles2 v.2.7.2 (Sedlazeck et al. 2018; Smolka et al. 2024), specifying the sample ID (*--sample-id*) and reference assembly (*--reference*) to allow for the output of sample-specific deletion sequences, in addition to the intermediate associated .snf file (*--snf*) necessary for efficient analyses at the cohort-level. Individual .snf files were then combined to jointly call structural variants in the two populations. Structural variant calls of different types (including deletions, duplications, insertions and inversions) were separated using BCFtools *view* v.1.22 (Danecek et al. 2021), keeping only those that passed the built-in quality criteria (*-f* PASS), and genotypes of insertions and deletions (indels) with a length of ≤ 10 kb were further refined using the *k*-mer genotyper kanpig (English et al. 2025) to improve their accuracy. In brief, using BCFtools *merge*, candidate indels ≤ 10 kb from each individual were first merged by ID (*-m* id) to avoid mis-merging of different variants at the same locus. Merged indels were left-aligned and normalized with regards to the reference assembly (*-f*), and multi-allelic sites split into biallelic sites (*-m -any*) using BCFtools *norm*. Truvari *collapse* version 5.4.0 (English et al. 2022) was then used to merge structural variant clusters and consolidate redundant calls (i.e., variants of the same type with ≥ 95% similarity in sequence and size, located within 500 bp), keeping only those with the highest quality score (*-k* maxqual) that passed built-in quality criteria (*--passonly*), and resolving symbolic variants with their sequence (*-w*) based on the information available in the reference assembly (*-f*). Afterward, variants observed in each individual (*--sample*) were re-genotyped using kanpig *gt* v.2.0.2 together with the long-read alignments (*--reads*) and reference assembly (*--reference*), keeping only those variants that passed built-in quality criteria (*--passonly*), and merged into a cohort-level dataset using BCFtools *merge*. BCFtools *filter* was used to remove any sites that were no longer segregating in the populations after re-genotyping as well as those for which the length reported for the structural variant (SVLEN) was inconsistent with the allelic sequence displayed for the alternate allele (ALT). Lastly, the cohort dataset was limited to high-confidence sites genotyped in all individuals. A summary of the structural variants discovered in the 20 individuals of this study is provided in Supplementary Table S3.

### The landscape of structural variation in rhesus macaques of Chinese and Indian descent

Population differentiation was assessed using Weir and Cockerham’s weighted *F_ST_* (1984) in R v.4.4.2 (R Core Team 2026). To test whether the relative composition of the structural variant landscape differed between the rhesus macaque populations of Chinese and Indian descent, a multivariate analysis was performed. To this end, the number of structural variants of each type were converted into proportions by normalizing by the total number of structural variants observed in an individual and a permutational multivariate analysis of variance was carried out using the *adonis2* function implemented in the *vegan* package (Oksanen et al. 2026), including the population as explanatory variable and assessing significance through permutation. Venn diagrams were plotted using the *ggVennDiagram* package (Gao et al. 2021). Population-specific differences in structural variant load and structural variant lengths were assessed using Wilcoxon rank-sum tests as implemented in R. Summary statistics were calculated for each structural variant type and each population separately, and resulting *p*-values were adjusted for multiple testing using the Benjamini-Hochberg method (2000). To ensure that observations were not driven by differences in sample sizes between the populations, statistical tests were repeated on balanced datasets generated by randomly sub-sampling variants from the more diverse (Chinese) population to match the sample size of the less diverse (Indian) population.

To test for the non-random distribution of structural variants across different chromosomes, observed per-chromosome structural variant counts in each population were compared to expectations under a null model assuming a uniform distribution proportional to the chromosome length. Deviations from expectation were assessed using a Poisson test, and *p*-values were adjusted for multiple testing using the Benjamini-Hochberg method (2000). Enrichments / depletions at the 5% significance level were considered significant. To assess whether the observed variation in structural variant density was associated with genomic features, a linear model was fit for each population and structural variant type, modelling structural variant density as a function of standardized (mean = 0, standard deviation = 1) measurements of three chromosomal features: recombination rate (based on the rates previously inferred by Versoza et al. 2024), GC-content, and repeat-content (both based on the annotations available for the rhesus macaque reference assembly; Warren et al. 2020). *P*-values were adjusted for multiple testing using the Benjamini-Hochberg method (2000) and results were visualized as coefficient plots of the effect sizes of each predictor. Additionally, to test for structural variant hotspots along each chromosome, structural variant counts of each type were divided into non-overlapping 1 Mb windows, normalized by bin size, and converted into *z*-scores based on chromosome- and structural variant type-specific distributions. Bins with *z*-scores ≥ 2 were considered hotspots of structural variation.

To predict functional consequences, structural variants were annotated using the Ensembl Variant Effect Predictor (VEP) v.115.2 (McLaren et al. 2016). To this end, a "cache" of the gene annotations for the rhesus macaque reference assembly, Mmul_10, containing information regarding species-specific transcript models and regulatory features, was downloaded (*vep_install -s* macaca_mulatta *-y* Mmul_10) and then used to annotate the population-scale variant catalogue (*vep --cache --species* macaca_mulatta), reporting predicted protein-level consequences (*--protein*) for variants up 200 Mb in size (*--max_sv_size* 200000000). Consequence terms reported by VEP were consolidated into biologically meaningful categories to reduce sparsity and improve interpretability. Specifically, structural variant consequences were grouped into six categories: protein-disrupting (consisting of frameshift variants, splice acceptor/donor variants, lost start/stop codons, transcript ablations, and feature truncations), coding changes (codon sequence/transcript variants), regulatory changes (downstream/upstream gene variants, 3’/5’ UTR variants, splice region variants, and feature elongations), non-coding transcript changes (intron variants, non-coding transcript variants, transcript amplification, and miRNA variants), intergenic changes (intergenic variants), and other (all remaining annotations). The distribution of both functional consequences and impacts predicted by VEP were visualized as bar plots in R containing the proportion of variants in each category, stratified by structural variant type and population. Differences in the distribution of functional consequences of deletions and insertions (i.e., the two variant types with sufficiently large sample sizes) between the two populations were assessed using χ^2^ tests.

### The landscape of mobile elements in rhesus macaques of Chinese and Indian descent

For each individual, autosomal mobile elements were identified from the long-read data using rMETL v.1.0.4 (Jiang et al. 2019). To this end, the *detection* function was first used to infer a dataset of candidate mobile element loci from the individual long-read alignments (*-x* pacbio) together with information from the reference assembly, requiring a minimum number of reads corresponding to half of an individual’s average read coverage to support a candidate (*-s*). Sections of reads identified as chimeric were then realigned based on known repetitive DNA elements in primates (as annotated in the Dfam database release 3.9; Storer et al. 2021) using the *realignment* function with the *-x* pacbio preset and a dataset of mobile elements was generated for each individual using the *calling* function. Annotations were limited to canonical mobile element classes (LINEs, SINEs, long terminal repeat retrotransposons, and DNA transposons) and population genetic analyses were limited to mobile elements with recent or current activity in primates (*Alu*, L1, and HERV-K(HML-2)-like elements).

To confirm population structure, a principal component analysis was performed based on segregating *Alu* elements and visualized by plotting the first two principal components in R v.4.4.2 (R Core Team 2026). Afterward, the total number of mobile elements per individual was calculated, and population-specific differences in both the number of mobile elements and the relative composition of mobile element classes were assessed using Wilcoxon rank-sum and χ^2^ tests, respectively. Finally, to quantify enrichment directionality, log-odds ratios were calculated. To this end, mobile element counts per class were converted into odds ratios using the Haldane-Anscombe correction, the variance of log-odds ratios was estimated using a Wald approximation, and statistical significance was assessed using *z*-tests. *P*-values were adjusted for multiple testing using the Benjamini-Hochberg method (2000).

### Inference of mutational mechanisms underlying structural variant formation

Long-read sequencing allows for the characterization of the precise breakpoints of structural variants, and patterns of microhomology in genomic regions flanking these breakpoints can be leveraged to infer the putative mutational mechanisms underlying the formation of these variants (Kidd et al. 2010; Lam et al. 2010). In order to obtain the precise sequences flanking the structural variant breakpoints, including any single nucleotide polymorphisms (SNPs) and short insertions / deletions (indels) segregating in individuals carrying the structural variants in question, non-structural variants were called in each individual using DeepVariant v.1.6.1 (Poplin et al. 2018) and genotyped jointly across the cohort using GLnexus v.1.4.1 (Yun et al. 2020) to improve genotyping accuracy. Afterward, cohort-level SNP, indel, and structural variant catalogues were limited to loci accessible in all individuals — defined here as genomic regions with a minimum depth of coverage of two in each individual (with the depth of coverage calculated using the *genomecov* function implemented in BEDTools v.2.30.0; Quinlan and Hall 2010) — split per individual (*--indv*) using VCFtools v.0.1.14 (Danecek et al. 2011), and then merged into a single catalogue per individual containing segregating variants of all types using BCFtools *concat* v.1.14 (Danecek et al. 2021). These complete variant catalogues per individual were phased using WhatsHap v.2.3 (Martin et al. 2016) and subsequently split by haplotype. Haplotype-resolved genome sequences of singletons were generated using BCFtools *consensus* together with the rhesus macaque reference assembly, Mmul_10 (GenBank accession number: GCA_003339765.3; Warren et al. 2020). Genomic regions flanking the breakpoints of non-mobile elements were extracted using BEDTools *getfasta*, limiting the analysis to structural variants farther than 200 bp from any other structural variant or indel on the same haplotype to avoid mis-inference. Following recent work in humans (Schloissnig et al. 2025), patterns of microhomology observed in these flanking regions were used to distinguish between structural variants originating from error in DNA double-strand repair via NHEJ (0 bp microhomologous sequences), MMEJ (1–15 bp), and NAHR (> 15 bp).

### Identification of population-private structural variants

To identify structural variants characterizing the Chinese and Indian populations, population-private structural variants were identified as those with a positive variant allele frequency in the focal population while being absent in the alternate population, requiring at least five individual genotype calls in each population. To assess statistical significance, the statistical framework of Maruki et al. (2022) was adapted for our structural variant calls. Specifically, the probability of a population-private structural variant call, *P*_*p*_, was calculated as *P*_*p*_ = 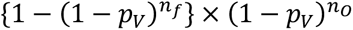, where *p_V_*, *n_F_*, and *n_O_* denote the total variant allele frequency in both populations, the number of allele calls in the focal population, and the number of allele calls in the alternate population, respectively. A Bonferroni correction was applied by dividing the critical value by the number of structural variants in variant allele frequency bins of size 0.05 to account for false positives resulting from multiple testing. Population-private structural variants at the 5% significance level were considered significant.

### Gene ontology enrichment analysis of population-private structural variants

To test whether certain biological functions or pathways were overrepresented among the population-private structural variants, gene ontology (GO) terms (Ashburner et al. 2000) of genes intersecting with population-private structural variants (as annotated in the rhesus macaque reference assembly, Mmul_10; Warren et al. 2020) were obtained using the DAVID web server (Huang et al. 2009a,b) and compared against a genomic background consisting of genes intersecting with all structural variants identified in the populations. Multiple testing correction was performed using the Benjamini-Hochberg procedure (2000), and GO terms with a false discovery rate < 0.05 were considered significant.

### Inference of diseases associated with significant population-private structural variants

To evaluate the potential biomedical relevance of the genetic differences observed between the rhesus macaque populations of Chinese and Indian descent, the association between significant population-private structural variants predicted to exhibit a high functional impact on genes (as annotated using VEP; see the section entitled "The landscape of structural variation in rhesus macaques of Chinese and Indian descent") and genes with a known disease-link in humans was examined using the eDGAR database (Babbi et al. 2017).

## ACKNOWLEDGEMENTS

We would like to thank Sam Peterson and the team at the Oregon National Primate Research Center (ONPRC) for providing the rhesus macaque samples used in this study. DNA extraction was performed at the ONPRC (Beaverton, OR, USA), library preparation and PacBio HiFi sequencing were performed at the Arizona Genomics Institute at the University of Arizona (Tucson, AZ, USA). Computations were performed on the Sol supercomputer at Arizona State University (Jennewein et al. 2023).

