## Supplementary Materials for "Evolutionary genomics based on PacBio HiFi long-read sequencing data reveals the importance of structural variants in shaping population-specific differences between Chinese and Indian rhesus macaques (*Macaca mulatta*)"

**Supplementary Table S1.** Sample information.

| | ID | read length (N50) | coverage |
|---|---|---|---|
| Chinese | RMac_01 | 16.8 kb | 10.9 |
| | RMac_02 | 21.4 kb | 6.3 |
| | RMac_03 | 22.7 kb | 14.5 |
| | RMac_04 | 22.9 kb | 8.0 |
| | RMac_05 | 23.2 kb | 12.5 |
| | RMac_06 | 22.0 kb | 16.2 |
| | RMac_07 | 22.9 kb | 12.7 |
| | RMac_08 | 23.2 kb | 12.4 |
| | RMac_09 | 18.2 kb | 10.5 |
| | RMac_10 | 22.3 kb | 13.3 |
| Indian | RMac_11 | 15.5 kb | 16.7 |
| | RMac_12 | 17.6 kb | 13.0 |
| | RMac_13 | 18.4 kb | 12.3 |
| | RMac_14 | 18.1 kb | 10.5 |
| | RMac_15 | 18.1 kb | 14.6 |
| | RMac_16 | 18.7 kb | 10.8 |
| | RMac_17 | 16.6 kb | 10.8 |
| | RMac_18 | 21.1 kb | 16.1 |
| | RMac_19 | 17.8 kb | 12.9 |
| | RMac_20 | 22.2 kb | 15.0 |

**Supplementary Table S2.** Summary of long-reads aligned to the Mmul_10 reference assembly.

| | ID | % aligned reads | alignment identity | mean mapping quality |
|---|---|---|---|---|
| Chinese | RMac_01 | 99.992 | 97.79 | 38.44 |
| | RMac_02 | 99.994 | 97.69 | 38.71 |
| | RMac_03 | 99.996 | 97.73 | 40.54 |
| | RMac_04 | 99.997 | 97.76 | 40.99 |
| | RMac_05 | 99.997 | 97.64 | 39.23 |
| | RMac_06 | 99.997 | 97.67 | 39.05 |
| | RMac_07 | 99.995 | 97.64 | 38.70 |
| | RMac_08 | 99.996 | 97.63 | 38.97 |
| | RMac_09 | 99.994 | 97.81 | 39.87 |
| | RMac_10 | 99.997 | 97.67 | 39.85 |
| Indian | RMac_11 | 99.993 | 98.19 | 38.33 |
| | RMac_12 | 99.995 | 98.09 | 38.38 |
| | RMac_13 | 99.995 | 98.05 | 37.80 |
| | RMac_14 | 99.993 | 98.04 | 37.81 |
| | RMac_15 | 99.995 | 98.07 | 38.22 |
| | RMac_16 | 99.994 | 98.03 | 37.75 |
| | RMac_17 | 99.994 | 98.13 | 38.34 |
| | RMac_18 | 99.996 | 97.97 | 38.10 |
| | RMac_19 | 99.993 | 98.10 | 38.69 |
| | RMac_20 | 99.996 | 97.98 | 38.24 |

**Supplementary Table S3.** Summary of structural variants.

| | ID | deletions | duplications | insertions | inversions |
|---|---|---|---|---|---|
| Chinese | RMac_01 | 37,822 | 11 | 55,955 | 80 |
| | RMac_02 | 34,971 | 6 | 49,177 | 57 |
| | RMac_03 | 38,414 | 12 | 57,257 | 86 |
| | RMac_04 | 36,267 | 9 | 52,367 | 67 |
| | RMac_05 | 37,875 | 11 | 56,490 | 75 |
| | RMac_06 | 38,392 | 10 | 58,005 | 88 |
| | RMac_07 | 37,899 | 9 | 56,395 | 78 |
| | RMac_08 | 36,234 | 15 | 53,213 | 90 |
| | RMac_09 | 37,009 | 11 | 54,332 | 79 |
| | RMac_10 | 38,468 | 7 | 57,503 | 77 |
| Indian | RMac_11 | 27,732 | 15 | 44,318 | 63 |
| | RMac_12 | 29,636 | 10 | 47,008 | 67 |
| | RMac_13 | 29,925 | 11 | 46,966 | 64 |
| | RMac_14 | 29,166 | 13 | 46,026 | 64 |
| | RMac_15 | 29,350 | 9 | 46,837 | 77 |
| | RMac_16 | 29,145 | 9 | 45,688 | 61 |
| | RMac_17 | 29,530 | 13 | 46,558 | 55 |
| | RMac_18 | 29,618 | 10 | 47,067 | 67 |
| | RMac_19 | 29,651 | 12 | 46,628 | 69 |
| | RMac_20 | 29,365 | 8 | 46,471 | 61 |

**Supplementary Table S4.** Gene ontology (GO) terms enriched in genes intersecting structural variants private to the Indian population. BP, CC, and MF denote the biological process, cellular component, and molecular function, respectively. Count and pop hits denote the numbers of genes associated with each term in the population-private structural variant catalogue and in the genomic background, respectively. FDR denotes the false discovery rate.

| category | term | count | pop hits | fold-enrichment | FDR |
|---|---|---|---|---|---|
| BP | GO:0006468~protein phosphorylation | 299 | 358 | 1.20 | 0.00001 |
| BP | GO:0007399~nervous system development | 215 | 252 | 1.22 | 0.00005 |
| BP | GO:0007156~homophilic cell-cell adhesion | 119 | 135 | 1.27 | 0.00283 |
| BP | GO:0006355~regulation of DNA-templated transcription | 463 | 591 | 1.12 | 0.00284 |
| BP | GO:0035556~intracellular signal transduction | 263 | 325 | 1.16 | 0.00684 |
| BP | GO:0007155~cell adhesion | 265 | 328 | 1.16 | 0.00684 |
| BP | GO:0007165~signal transduction | 368 | 471 | 1.12 | 0.03141 |
| BP | GO:0007420~brain development | 104 | 120 | 1.24 | 0.03141 |
| CC | GO:0098978~glutamatergic synapse | 290 | 352 | 1.19 | 0.00005 |
| CC | GO:0045211~postsynaptic membrane | 131 | 148 | 1.27 | 0.00009 |
| CC | GO:0005829~cytosol | 2,136 | 2,922 | 1.05 | 0.00045 |
| CC | GO:0030424~axon | 182 | 217 | 1.21 | 0.00061 |
| CC | GO:0005886~plasma membrane | 2,271 | 3,123 | 1.05 | 0.00126 |
| CC | GO:0043197~dendritic spine | 75 | 82 | 1.32 | 0.00165 |
| CC | GO:0098794~postsynapse | 104 | 120 | 1.25 | 0.00528 |
| CC | GO:0045202~synapse | 257 | 325 | 1.14 | 0.01783 |
| CC | GO:0005737~cytoplasm | 2,785 | 3,880 | 1.03 | 0.02243 |
| CC | GO:0098685~Schaffer | 60 | 67 | 1.29 | 0.04711 |
| MF | GO:0005524~ATP binding | 1,179 | 1,293 | 1.09 | 0.00000 |
| MF | GO:0005515~protein binding | 1,707 | 1,925 | 1.06 | 0.00000 |
| MF | GO:0005509~calcium ion binding | 509 | 564 | 1.08 | 0.01397 |
| MF | GO:0031267~small GTPase binding | 193 | 206 | 1.12 | 0.04297 |

**Supplementary Table S5.** Gene ontology (GO) terms enriched in genes intersecting structural variants private to the Chinese population. BP, CC, and MF denote the biological process, cellular component, and molecular function, respectively. Count and pop hits denote the numbers of genes associated with each term in the population-private structural variant catalogue and in the genomic background, respectively. FDR denotes the false discovery rate.

| category | term | count | pop hits | fold-enrichment | FDR |
|---|---|---|---|---|---|
| BP | GO:0007156~homophilic cell-cell adhesion | 132 | 135 | 1.17 | 0.00444 |
| BP | GO:0006468~protein phosphorylation | 331 | 358 | 1.11 | 0.00444 |
| BP | GO:0035556~intracellular signal transduction | 300 | 325 | 1.10 | 0.01556 |
| BP | GO:0006897~endocytosis | 146 | 153 | 1.14 | 0.04751 |
| CC | GO:0005829~cytosol | 2,549 | 2,922 | 1.04 | 0.00000 |
| CC | GO:0005737~cytoplasm | 3,351 | 3,880 | 1.03 | 0.00001 |
| CC | GO:0005856~cytoskeleton | 357 | 389 | 1.10 | 0.00151 |
| CC | GO:0005925~focal adhesion | 155 | 162 | 1.14 | 0.00255 |
| CC | GO:0005654~nucleoplasm | 1,750 | 2,019 | 1.04 | 0.00845 |
| CC | GO:0005929~cilium | 178 | 190 | 1.12 | 0.01715 |
| CC | GO:0015629~actin cytoskeleton | 175 | 187 | 1.12 | 0.02060 |
| CC | GO:0042995~cell projection | 141 | 149 | 1.13 | 0.02261 |
| CC | GO:0030424~axon | 201 | 217 | 1.11 | 0.02546 |
| CC | GO:0098978~glutamatergic synapse | 319 | 352 | 1.08 | 0.02546 |
| CC | GO:0005912~adherens junction | 120 | 126 | 1.14 | 0.02779 |
| CC | GO:0016604~nuclear body | 223 | 243 | 1.10 | 0.03798 |
| MF | GO:0005524~ATP binding | 1,179 | 1,293 | 1.09 | 0.00000 |
| MF | GO:0005515~protein binding | 1,707 | 1,925 | 1.06 | 0.00000 |
| MF | GO:0005509~calcium ion binding | 509 | 564 | 1.08 | 0.01397 |
| MF | GO:0031267~small GTPase binding | 193 | 206 | 1.12 | 0.04297 |

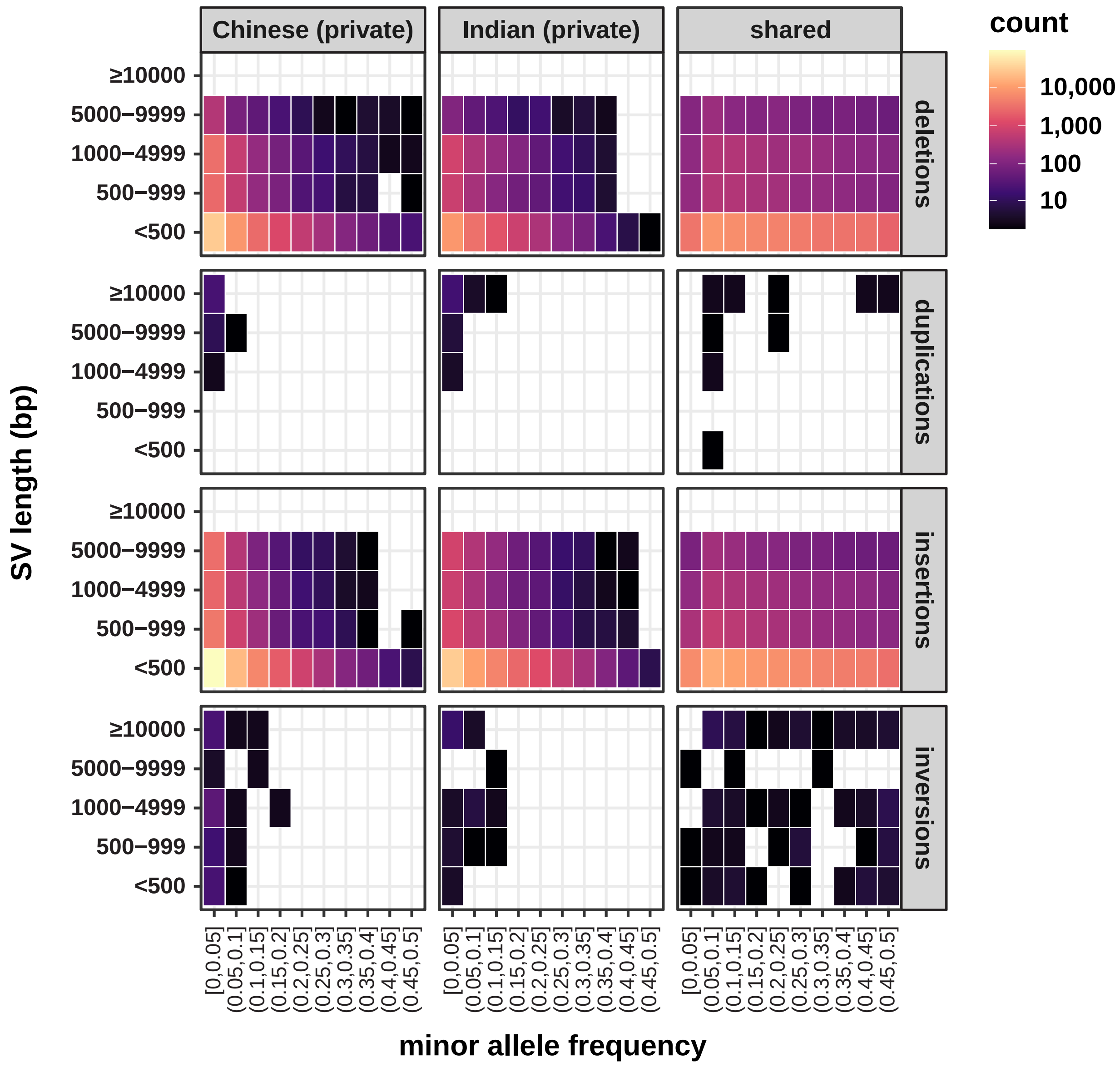


**Supplementary Figure S1.** Minor allele frequency of structural variants (deletions, duplications, insertions and inversions) private to the Chinese population (left panel), private to the Indian population (middle panel), and shared between the two populations (right panel) stratified by their length.

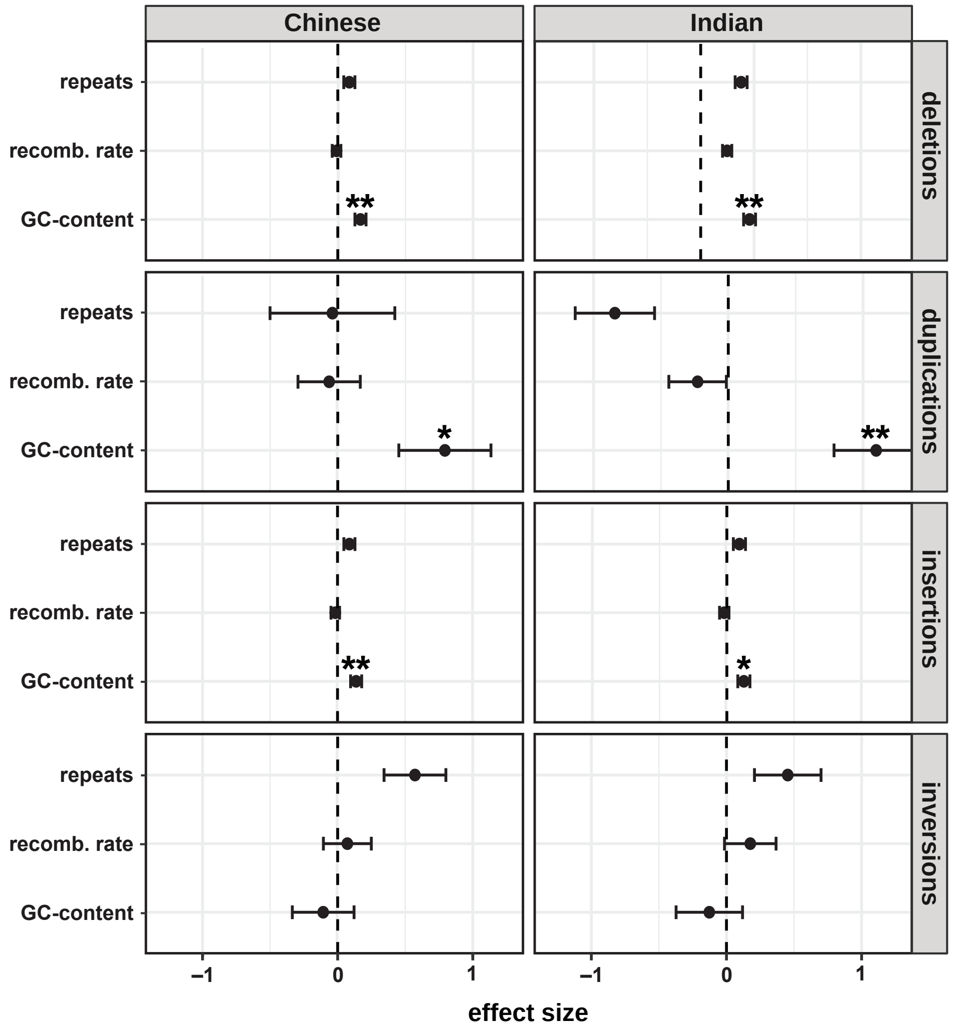


**Supplementary Figure S2.** Coefficient plots of the effect sizes of recombination rate (based on the rates previously inferred by Versoza et al. 2024), GC-content, and repeat-content (both based on the annotations available for the rhesus macaque reference assembly; Warren et al. 2020) as predictors of structural variant density at the chromosomal scale. * and ** indicate a *p*-value < 0.05 and < 0.01, respectively.

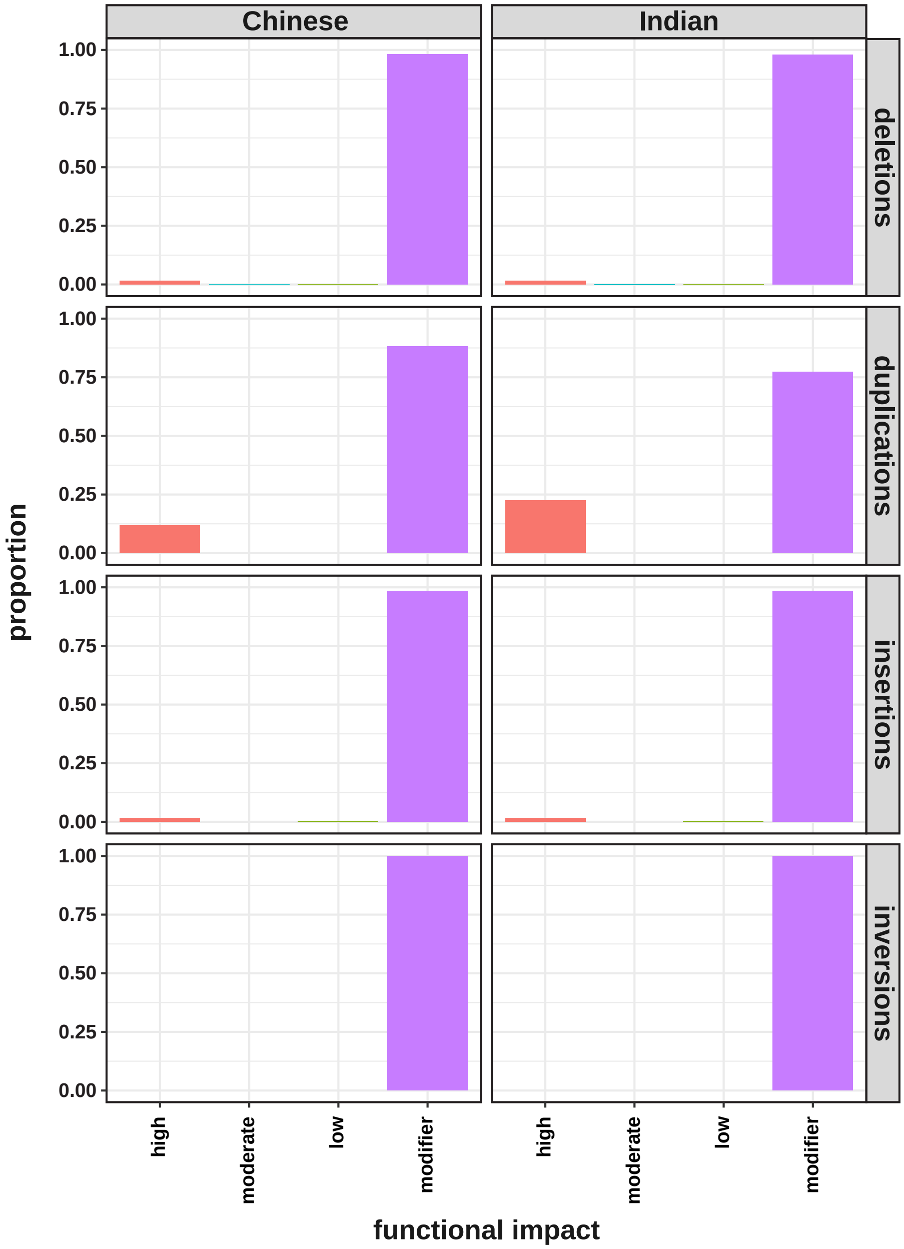


**Supplementary Figure S3.** Distribution of population-specific structural variants across four functional categories of decreasing impact: high, moderate, and low effect, as well as modifiers with little to no predicted effect.
